# Identifying factors that contribute to collision avoidance behaviours while walking in a natural environment

**DOI:** 10.1101/2024.06.11.598509

**Authors:** Mohammadamin Nikmanesh, Michael E. Cinelli, Daniel S. Marigold

## Abstract

Busy walking paths, like in a park, a sidewalk in a city centre, or a shopping mall, frequently necessitate collision avoidance behaviour. Lab-based research has shown how a variety of situation-specific factors (e.g., distraction, object/pedestrian proximity) and person-specific factors (e.g., pedestrian size, age), typically studied independently, affect avoidance behaviour. What happens in the real world is unclear. Thus, we filmed unscripted pedestrian walking behaviours on a busy ∼3.5 m urban path adjacent to the water. We leveraged deep learning algorithms to identify and extract walking trajectories of pedestrians and had unbiased raters characterize interaction details. Here we analyzed over 500 situations where two pedestrians approached each other from opposite ends (i.e., one-on-one pedestrian interactions). We found that smaller medial-lateral distance between approaching pedestrians and a lower number of surrounding pedestrians (i.e., smaller crowd size) predicted an increase in the likelihood of a subsequent path deviation. Furthermore, we found that whether a pedestrian looked distracted or held, pushed, or pulled something while walking predicted the medial-lateral distance between pedestrians at the time of crossing. Although pedestrians maintained a larger personal space boundary compared to lab settings, this is likely because of the outdoor path’s width. Overall, our results suggest that collision avoidance behaviours in lab and real-world environments share similarities and offer insights relevant to developing more accurate computational models for realistic pedestrian movement.

## INTRODUCTION

Navigation is fundamental to our daily lives. Consider the common experience of walking along a busy path in a park or through a crowded mall. In each case, a person is challenged to successfully negotiate a space shared with others, circumventing individual and groups of pedestrian(s) and physical barriers, like a bench or a kiosk. Decision-making while navigating crowded spaces involves adjusting to an ever-changing environment. But how are decisions made about which path to walk? And more specifically, which factors influence a pedestrian’s decision to deviate from their current trajectory, independent of whether a collision is probable?

Research suggests that situation-specific factors, like intended heading direction, walking speed, obstacle/pedestrian proximity and orientation, crowd density, distracted pedestrians, and path width, affect collision avoidance behaviour (Cutting et al. 1995; Huber et al. 2014; Gérin-Lajoie et al. 2006; Knorr et al. 2016; Meerhoff et al. 2018; Rapos et al. 2019, 2021). For example, avoiding a collision with an object or person approaching along a 180-degree path (head on) requires a change in pathway, whereas avoiding an object or person approaching along a 90-degree path requires a change in speed. Furthermore, pedestrians increase their trajectory deviation when passing someone distracted by using a mobile phone (Murakami et al. 2022). However, observable personal characteristics of approaching pedestrians (e.g., age, body size) also have the potential to alter locomotor trajectory and avoidance behaviours.

There are relatively fewer studies investigating the role of person-specific factors on human collision avoidance behaviour. Bourgaize et al. (2021) found that body size (height, weight, and shoulder width) of an interferer affected medial-lateral clearance at the time of crossing, such that a larger interferer induced a greater clearance. In addition, people prefer to intrude the personal space of short pedestrians compared to tall pedestrians (Caplan and Goldman 1981). Age of an approaching pedestrian also appears to affect one’s avoidance behaviours. Bourgaize et al. (2023) found that when an approaching pedestrian looked like an older adult or walked like an older adult, young adults increased their medial-lateral clearance at the time of crossing. These findings highlight the importance of considering a wide range of factors in understanding and predicting pedestrian collision avoidance behaviour.

Computational models that help explain how people select locomotor trajectories and avoid collisions tend to ignore person-specific characteristics. Most models that explain collision avoidance behaviours tend to treat other pedestrians and other obstacles in an environment as points in space (Fajen 2013; Fajen and Warren 2003; Helbing and Molnar 1995). For instance, the Behavioral Dynamics model (Fajen and Warren 2003) treats objects within an environment as repellers (causing one to deviate from their path) and goals as attractors (used to set up locomotor trajectory). The magnitude of repulsion or attraction within this model is based on one’s proximity to the obstacle or goal respectively, regardless of their properties. The Behavioral Dynamics Model does a fairly good job of reproducing avoidance behaviours in laboratory settings with stationary and moving objects. Similarly, the Social Force model (Helbing and Molnar 1995) reproduces pedestrian interactions with the driving force of the intended destination, repulsive forces of other pedestrians and obstacles, and attraction forces of friends or other individuals of interest. The latter model, and its many revisions, are often used to understand crowd behaviour, including simulations of building evacuation (Chen et al. 2018; Farina et al. 2017).

Our understanding of individuals’ behaviours has primarily emerged from controlled laboratory settings, where factors that affect collision avoidance are examined in isolation. While laboratory-based studies provide clarity on these different factors, they fail to account for their combined effect in real-life environments. In real-life, pedestrians are constantly perceiving environmental, personal, and situation-specific factors simultaneously. To bridge the gap between controlled laboratory settings and real-world conditions, we assessed unscripted pedestrians’ avoidance behaviours while walking along a paved path in a busy urban setting. We filmed pedestrian interactions to capture natural behaviour. From all the video footage we recorded, we focused our analysis here on situations in which there were only two pedestrians approaching each other from opposite ends of a path (i.e., one-on-one pedestrian interactions). We were interested in determining which factors—crowd size, medial-lateral distance from each other, age of pedestrians, presence of an obstacle, and whether one looked distracted or had their mobility constrained due to holding, pushing, or pulling an object or animal—predicted a pedestrian’s decision to deviate from their initial walking trajectory. We also sought to determine which of these factors predicted medial-lateral separation between pedestrians at the time of crossing. To help address our aims, we leveraged deep learning algorithms to identify and extract the locomotor trajectories of each pedestrian.

## METHODS

To address our research questions, we filmed natural walking behaviours along a paved path and used deep learning computer algorithms and subjective observation to analyze pedestrian interactions. The steps are illustrated in Figure 1.

**Figure 1.**
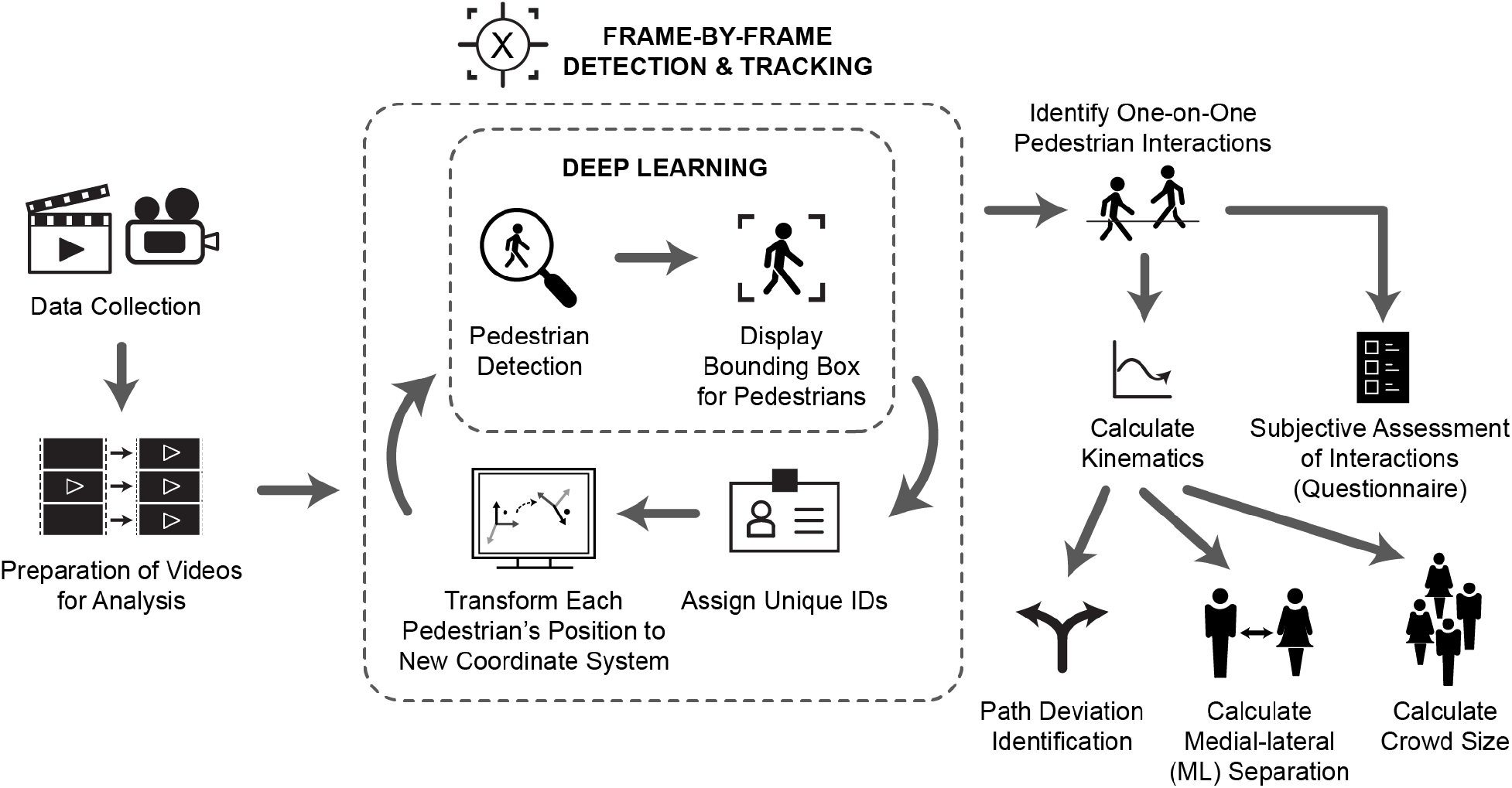
Illustration of the steps involved in data collection and analysis.

### Data collection

We filmed pedestrians walking along a 3.5 – 3.8 m wide urban path at English Bay, Vancouver, BC, Canada. A Sony FDR-AX43 UHD 4K Handycam Camcorder secured on a tripod recorded behaviour on an angle from an elevated position overlooking the path at 60 Hz. This elevated angle ensured the pedestrian tracking algorithms would work. We selected this path for its high foot-traffic and location. This location is a confined path with a minimum number of obstacles. We collected data between the hours of noon to 5 pm during the peak pedestrian traffic months of June, July, and August of 2020 and 2021. The consistency in the time and month of data collection helped maintain a uniform pattern in pedestrian activity, contributing to the accuracy of our findings. Pedestrians were not aware of being filmed and thus, we were able to collect natural, real-life behaviour.

Because we filmed at a public place, there was no reasonable expectation of privacy. In addition, the research team did not interact with any pedestrian, nor collect any personal data. Thus, we did not require consent. The Office of Research Ethics at Simon Fraser University approved the study protocol.

### Preparation of videos for analysis

To make the kinematic analysis easier and more intuitive, we transformed videos into a bird’s-eye view (described in more detail later). Specifically, we created a perspective transformation matrix in Python to use later during the video analysis. We selected four points, forming a rectangular area of interest (AOI), from the first frame of the video that served as inputs to this function. We used the bottom left corner of the AOI as the origin of the new coordinate system (0, 0). Subsequently, we selected three points with a known distance between each to use for converting pixels to cm during the video analysis. Finally, we selected a point on the bottom left corner of any obstacles present (e.g., bench, garbage can, light pole).

### Frame-by-frame tracking: Identification of pedestrians and one-on-one pedestrian interactions

To detect pedestrians for further analysis, we processed the raw video footage using a deep learning algorithm called FairMOT, which is based on the YOLOV5 architecture (Zhang et al. 2021). FairMOT is specifically designed to identify pedestrians in videos and track their movements throughout the recorded period. We used a pre-trained version of this algorithm and customized it to suit the requirements of our project. This customization included use of this algorithm and embedded pedestrian tracking, converting distances, and extracting kinematic data. One of the key capabilities of FairMOT is its ability to track multiple pedestrians simultaneously, allowing us to extract each pedestrian’s individual walking trajectory. The algorithm analyzes frames from the video using an encoder-decoder network. With this network, the encoder compresses the input, and the decoder reconstructs features relevant to pedestrian detection. This generates a feature map that highlights elements indicative of human presence, allowing for efficient detection and tracking of pedestrians.

Each frame, all the pixels that make up pedestrian(s) are identified, then the algorithm creates a distinct bounding box, with a unique ID, around each separate pedestrian. This bounding box not only serves to distinguish between individuals but also provides a means to measure distances between them. Subsequently, we transform the lower left corner of each bounding box to a bird’s-eye view using the perspective transformation matrix created earlier. Finally, for each point representing a bounding box, we convert the pixel coordinates to cm. To perform a pixel-to-cm conversion, we used following equations:

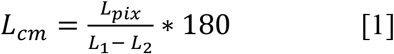

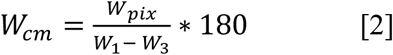

where *L1* – *L2* represents the difference in pixel length (y-coordinate) between two of the selected points that correspond to 180 cm in the real world. Similarly, *W1* – *W3* represents the difference in pixel width (x-coordinate) between two of the selected points that correspond to 180 cm in the real world. *L*_*pix*_ and *W*_*pix*_ represent the position in pixels of a point in the video and *L*cm and *W*cm represent the position in cm of that point.

To track each pedestrian across frames, we used *JDETracker* from the FairMOT library. This method compares the locations of pedestrians in consecutive frames. If the coordinates of an individual (i.e., the point representing the bounding box) in the current frame are within a specified proximity to the coordinates of the same person in the previous frame, they are considered as from the same pedestrian.

After detecting and tracking all pedestrians in the video, we manually searched for one-on-one pedestrian interactions. We use the term interaction to denote an occurrence where one pedestrian walks past another pedestrian on the path; we did not observe any collisions. For each interaction, we recorded the pedestrian IDs and the time at which they walked past each other (i.e., crossing time) for later analyses.

### Path deviation identification

To address our primary question as to which factors influence whether a person deviates from their initial walking trajectory, we first had to determine whether and when one or both pedestrians deviated. We used a change in the medial-lateral (ML) separation between the two pedestrians, which represents the mutual ML separation. Specifically, we calculated the mean and standard deviation of the ML separation between the two pedestrians from 1 s before to 0.5 s after the crossing time. We then worked backwards in time from the time that the two pedestrians crossed each other to detect a point of deviation. A point of deviation was identified if the ML separation between the two pedestrians fell below 2 standard deviations of the average ML separation. This approach allowed us to identify the closest time to the time of crossing when two pedestrians increased their ML separation to avoid a collision while approaching each other. This method ensured that we detected the point of deviation related to the pedestrians of interest rather than deviations that might have occurred earlier due to interactions with other individuals.

### Calculation of crowd size

All deviations occurred when the two pedestrians were less than 8 m apart in the anterior-posterior direction (see Results). Thus, we used an 8 m distance as a reference point for calculating crowd size. To calculate the crowd size, we identified the precise time when two pedestrians were 8 meters apart from each other. At this moment, we counted the number of pedestrians within the designated scene. The designated scene included the 8 m distance between pedestrians, plus an additional 2 m behind each pedestrian, resulting in a total of 12 m in the anterior-posterior direction. In the ML direction, the designated scene included the ML distance between the two pedestrians, plus an additional 0.8 m on the outer side of each pedestrian. We included the extra area around each pedestrian because it provided sufficient space to include pedestrians who were slightly outside the direct line between the two individuals but still within a reasonable proximity who could theoretically influence deviation behaviour. By defining the scene dimensions in this manner, we ensured that our crowd size calculations captured a comprehensive view of the surrounding pedestrians involved in the interaction.

### Subjective assessment of interactions

We recruited people not part of the research team to review videos and answer a comprehensive questionnaire of 14 questions related to each one-on-one pedestrian interaction. We used three independent people for each interaction to ensure unbiased responses. The uneven number of people ensured that we had a tiebreaker in cases where two people disagreed on an answer. The questionnaire captured aspects of pedestrian interactions that could not be quantified merely through our kinematic data, such as the approximate age of the pedestrians, potential mobility constraints, distractions like phone usage, and reasons for deviating from a straight line even when far from another person. Each person rated their confidence level in their answers on a scale from 0 to 100%. We only used responses in later analyses where the average confidence level across all people exceeded 75%.

Some of the questionnaire questions had a greater number of categories than we opted to use in our analyses. We decided to use many categories to provide a thorough account of each interaction. However, to increase counts in certain categories and simplify analyses, we reduced the number of categories. Specifically, we reduced the age variable to two categories: pedestrians within the same age group and pedestrians of different age groups. Age groups included children (e.g., < 12 years old), adolescent or young adult (e.g., 13 – 20 years old), early adulthood (e.g., 21 to 40 years old), middle adulthood (e.g., 41 to 60 years old), and older adult (e.g., 60+ years old). Although we asked about a variety of different obstacles within a distance to affect behaviour (e.g., light pole, bench, stationary person), we used two categories for the presence of an obstacle: yes and no. In addition, we reduced the mobility constraint variable to two categories: yes and no. We defined a mobility constraint as an object held or used while walking, excluding a cell phone, and options on the questionnaire included a cane, two- or four-wheel walker, pushing a bicycle, stroller, or wheelchair, and walking an animal. Due to the challenge of identifying sex from the videos based on feedback from the people completing the questionnaire, we chose to remove this variable from further analyses. <u>Table</u> provides an overview of the factors we used in statistical analyses.

### Statistical analyses

To determine which factors can predict whether one or both pedestrians deviate during their interaction, we used a multiple logistic regression model. We included the binary response variable, deviation (yes/no; reference = no deviation), and the following predictor variables: ML separation at 8 m, crowd size at 8 m, age group, distraction, presence of obstacles, and mobility constraint.

In a series of secondary analyses, we first focussed only on interactions with deviations to determine which factors can predict the time of path deviation relative to the pedestrians walking past each other (i.e., time to pass from deviation). In a linear regression model, we included time to pass from deviation as the response variable (log transformed to ensure normality) and ML separation at 8 m, crowd size at 8 m, age group, distraction, and mobility constraints as predictor variables. We excluded the obstacle variable in this analysis because there were too few cases. Subsequently, we sought to determine which factors can predict the ML separation between the two pedestrians at the time of crossing (including cases with and without path deviations). We used a linear regression model, with ML separation (at time of crossing) as the response variable and age group, distraction, mobility constraints, presence of obstacles, and crowd size at 8 m distance between pedestrians as predictor variables.

We used RStudio version 2023.06.1+524 (R version 4.3.1) and an alpha level of 0.05 for all statistical analyses. We used the plot_model function of the sjPlot package version 2.8.15 to create plots of the results. We also re-run each regression model with only the significant predictors and show these results alongside the full model.

### Data and code availability

We will make data and code available in a repository at the time of publication.

## RESULTS

We identified a total of 521 one-on-one pedestrian interactions in our dataset. We excluded 79 interactions because either: (1) one or more of the pedestrians involved in the interaction were not in the video frame for enough time to gather sufficient kinematic data; (2) the answers on the questionnaires from the three volunteers were not consistent with one another; or (3) the confidence level in the answers provided by the volunteers was less than 75%. In the end, we analyzed 442 interactions, of which, one or both pedestrians deviated from their walking trajectory in 193 (44%) cases. All deviations occurred when pedestrians were between 1 and 8 m apart in the direction of walking, with 76% of cases occurring between 2 and 5.5 m apart (Figure 2).

**Figure 2.**
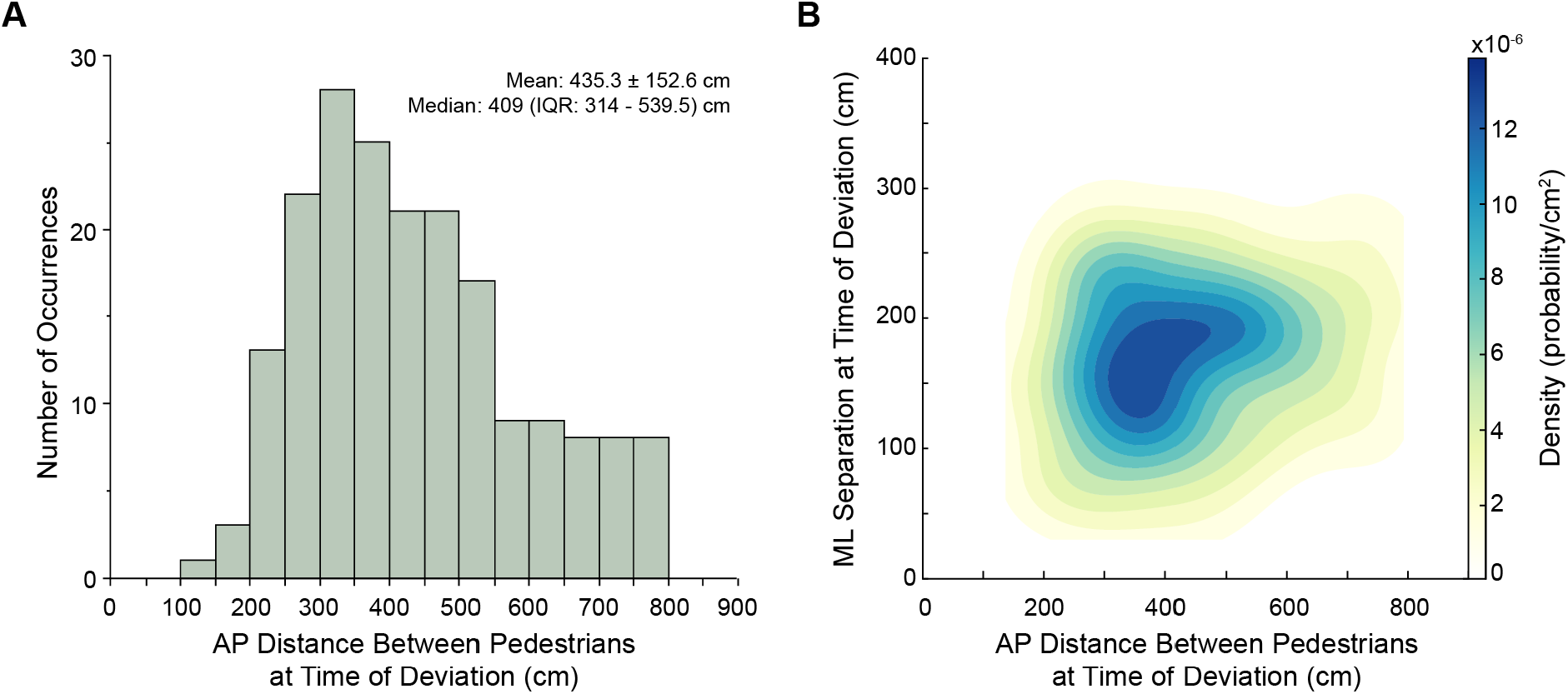
Distance between pedestrians at time of path deviation. **A:** Histogram of anterior-posterior (AP) distance between pedestrians. **B:** Density plot showing the relationship between ML separation and AP distance between pedestrians for cases where a path deviation occurred.

### Factors predicting path deviation

We ran a multiple logistic regression to determine which factors best predicted whether a deviation would occur. The logistic regression model included: (1) ML separation at 8 m apart; (2) crowd size at 8 m apart; (3) whether pedestrians were of the same age group; (4) whether pedestrians were distracted; (5) whether pedestrians had their mobility constrained due to holding, pushing, or pulling an object or animal; and (6) whether an obstacle (e.g., bench, light pole, stationary person) was present within a distance to affect behaviour. Both ML separation and crowd size significantly predicted path deviation (Table 2; Figure 3A,B). Specifically, we found an increase in the probability of deviation with a decrease in ML separation between pedestrians at 8 m apart (Figure 3C). This probability decreased for a given ML separation with larger crowd size. Together, these results highlight the importance of spatial considerations on path deviations.

**Table 1.**
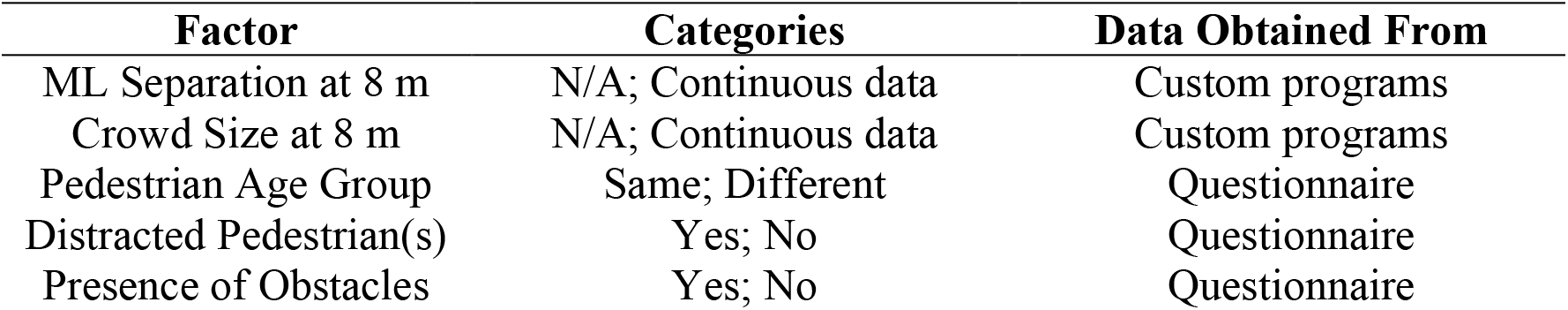
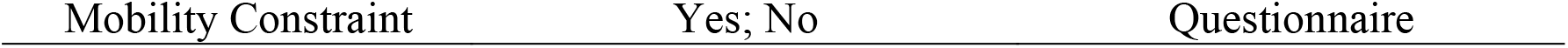
Factors that may influence path deviation and their categories.

**Table 2.** Results of the multiple logistic regression for factors predicting path deviation.

| Variable | Coefficient<br>($\beta$) | SE | $\chi^2$ | P-value | Odds Ratio | 95% CI |
| --- | --- | --- | --- | --- | --- | --- |
| (Intercept) | 2.196 | 0.373 | - | - | 8.988 | 4.422, 19.147 |
| ML separation | -0.014 | 0.002 | 76.608 | **< 2e-16** | 0.986 | 0.982, 0.989 |
| Age(Same) | 0.224 | 0.220 | 1.044 | 0.307 | 1.251 | 0.814, 1.930 |
| Distraction(Yes) | 0.402 | 0.317 | 1.603 | 0.205 | 1.495 | 0.801, 2.792 |
| Mobility<br>Constraint(Yes) | -0.241 | 0.306 | 0.628 | 0.428 | 0.786 | 0.428, 1.424 |
| Obstacle(Yes) | 0.605 | 0.502 | 1.431 | 0.232 | 1.830 | 0.674, 4.925 |
| Crowd # | -0.417 | 0.195 | 4.98 | **0.026** | 0.659 | 0.442, 0.952 |

**Figure 3.**
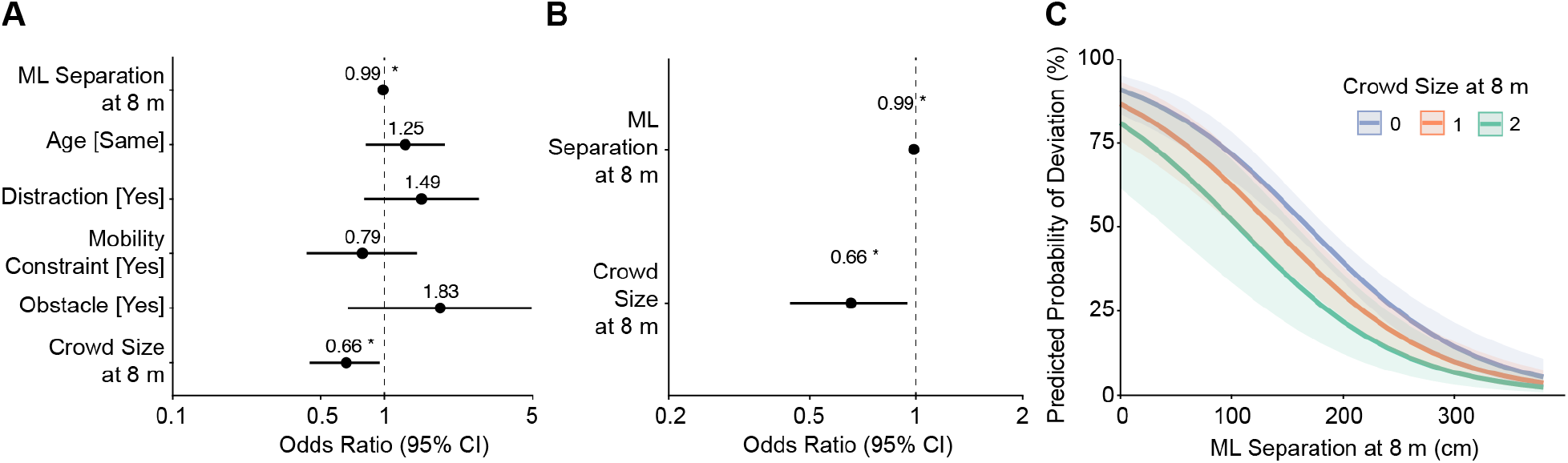
Factors predicting path deviation. **A:** Odds ratios of the factors in the full multiple logistic regression model. **B:** Odds ratios of the factors in the reduced multiple logistic regression model. **C**. Predicted probability of path deviation based on crowd size and ML separation at 8 m. Asterisk indicates a significant predictor variable (p < 0.05).

### Factors predicting the time for pedestrians to pass each other from the moment of deviation

Amongst cases where one or both pedestrians deviated from their original walking trajectory, we asked which factors predicted the time to pass from the moment of deviation. Factors in the multiple linear regression included: (1) ML separation at 8 m apart; (2) crowd size at 8 m apart; (3) whether pedestrians were of the same age group; (4) whether pedestrians were distracted; and (5) whether pedestrians had their mobility constrained due to holding, pushing, or pulling an object or animal. Only the ML separation when pedestrians were 8 m apart predicted the time to pass from deviation (Table 3; Fig. 4A,B). Time to pass from deviation decreased with decreasing ML separation (Fig. 4C).

**Table 3.** Results of the multiple linear regression for factors predicting the time to pass from deviation.

| Variable | Coefficient ($\beta$) | 95 % CI | P-value |
| --- | --- | --- | --- |
| (Intercept) | 0.119 | -0.019, 0.258 | 0.091 |
| ML separation | 0.001 | 0.001, 0.002 | **0.0003** |
| Crowd # | -0.033 | -0.137, 0.072 | 0.538 |
| Mobility Constraint(Yes) | 0.066 | -0.081, 0.212 | 0.378 |
| Age(Same) | -0.079 | -0.184, 0.026 | 0.139 |
| Distraction(Yes) | 0.015 | -0.128, 0.159 | 0.835 |

**Figure 4.**
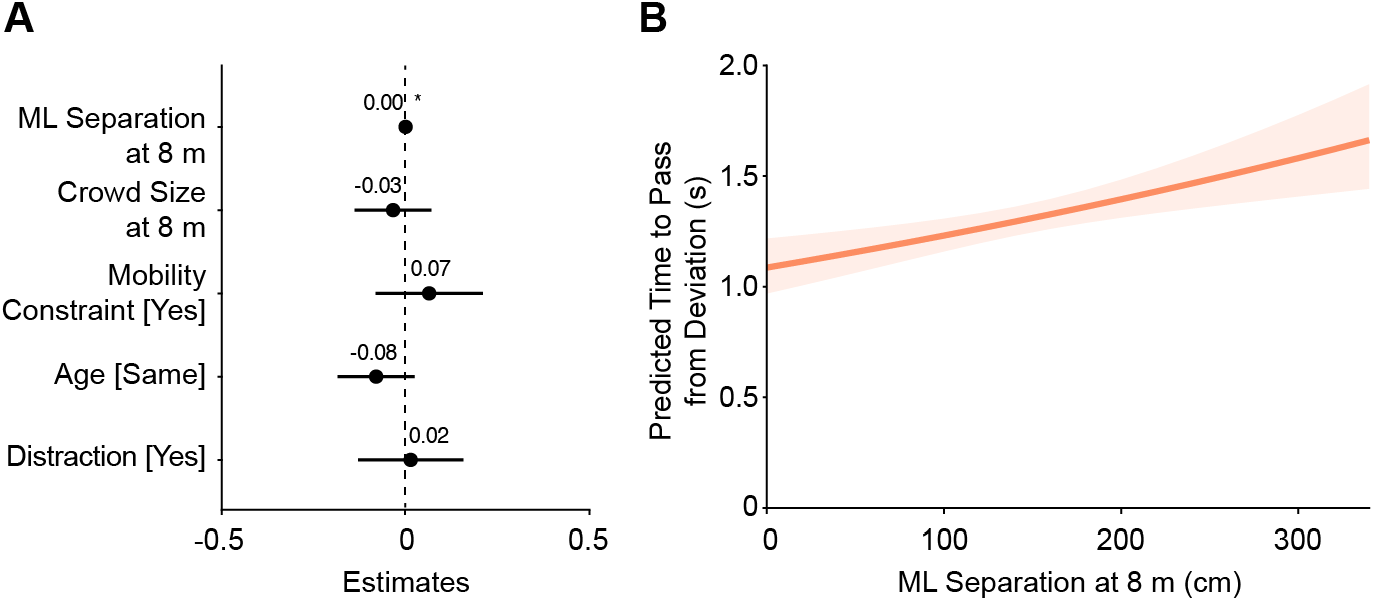
Factors predicting time to pass from deviation. **A:** Estimates of the factors in the full multiple linear regression model. Reduced model not shown because only one factor (ML separation at 8 m) is significant. **B:** Predicted time to pass from deviation based on ML separation at 8 m. Asterisk indicates a significant predictor variable (p < 0.05).

### Factors predicting ML separation at the time of crossing

Here we asked which factors predict ML separation at the time of crossing, including: (1) crowd size at 8 m apart; (2) whether pedestrians were of the same age group; (3) whether pedestrians were distracted; (4) whether pedestrians had their mobility constrained due to holding, pushing, or pulling an object or animal; and (5) whether an obstacle was present. Both mobility constraint and distraction significantly predicted ML separation at the time of crossing (Table 4; Fig. 5A,B). We found decreased ML separation with the presence of a mobility constraint (Fig. 5C), and increased ML separation when at least one of the pedestrians was distracted (Fig. 5C).

**Table 4.** Results of the multiple linear regression for factors predicting ML separation at time of crossing.

| Variable | Coefficient ($\beta$) | 95 % CI | P-value |
| --- | --- | --- | --- |
| (Intercept) | 185.033 | 175.472, 194.593 | $< 2e-16$ |
| Crowd # | 3.965 | -4.850, 12.779 | 0.377 |
| Mobility Constraint(Yes) | -23.811 | -39.548, -8.073 | **0.003** |
| Obstacle(Yes) | 5.038 | -20.731, 30.806 | 0.701 |
| Age(Yes) | 5.509 | -5.663, 16.682 | 0.333 |
| Distraction(Yes) | 21.478 | 5.450, 37.506 | **0.009** |

**Figure 5.**
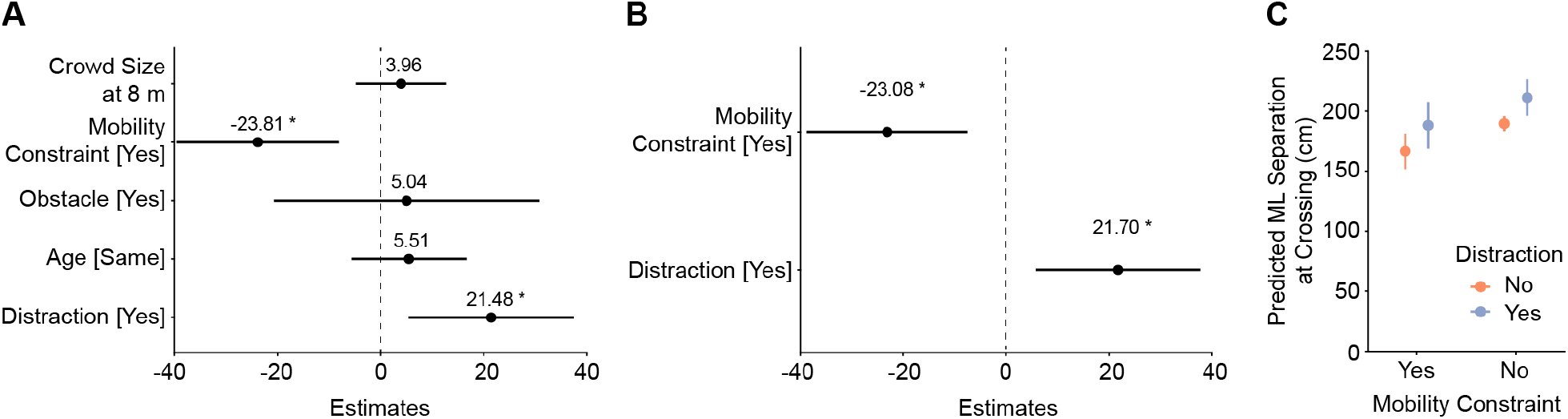
Factors predicting ML separation at time of pedestrian crossing. **A:** Estimates of the factors in the full multiple linear regression model. **B:** Estimates of the factors in the reduced multiple linear regression model. **C**. Predicted ML separation at crossing based on whether pedestrians were distracted and had a mobility constraint. Asterisk indicates a significant predictor variable (p < 0.05).

## DISCUSSION

Busy walking paths, like in a park, downtown sidewalk, or mall, commonly necessitate collision avoidance behaviour. In the current study, we determined which factors—crowd size, distance from each other, age, presence of obstacles, and whether one pedestrian looked distracted or had their mobility constrained due to holding, pushing, or pulling an object or animal—predicted a pedestrian’s decision to deviate from their initial walking trajectory. We also determined which of the above-mentioned factors predicted ML separation between pedestrians at the time of crossing. We focused on unscripted one-on-one pedestrian interactions based on video from a real-world public path and thus, completely natural behaviour. Our results revealed that ML separation and crowd size when pedestrians were 8 m apart predicted a subsequent path deviation, and the former also predicted the time to pass from the deviation point. Our results also revealed that whether a pedestrian looked distracted or had their mobility constrained predicted the ML separation at the time of crossing. These findings highlight the rich insight that observing natural pedestrian behaviour provides.

The likelihood of path deviation increased as ML separation between pedestrians at 8 m apart decreased. We found a 50% probability of path deviation when ML separation at 8 m was ∼175 cm (with no other pedestrians in the vicinity, i.e., crowd size = 0 in Figure 3C). Since each pedestrian’s position is tracked based on the lower left corner of the bounding box in the video, we must account for the shoulder width of one of the pedestrians to approximate the true ML separation. Assuming an average shoulder width of 45 cm, this equates to 130 cm ML separation. We found a 75% probability of path deviation when ML separation at 8 m was ∼90 cm (or 45 cm when accounting for the bounding box). With very small ML separation values, when a collision is imminent, it is reasonable to expect path deviations to occur. However, path deviations frequently occurred even in the absence of a probable collision. What might explain this behaviour? One possible reason is that pedestrians tried to maintain their personal space envelope as they walked past each other (Bourgaize et al. 2021; Gerin-Lajoie et al. 2005; Gorrini et al. 2014; Hecht et al. 2019; Liu et al. 2019; Shimizu et al. 2020). During locomotion, the personal space envelope is a protective zone around body that allows the person to detect hazards and initiate timely avoidance maneuvers, if necessary (Gerin-Lajoie et al. 2005). The typical ML separation distance at the time of crossing when circumventing a person/mannequin is approximately 50 – 70 cm (Bourgaize et al. 2021; Gerin-Lajoie et al. 2005; Shimizu et al. 2020). These values are based on the distance between the arm of one person/mannequin and the centre of mass of the other, or the distance between to the sternum of each person. Thus, when considering only the distance between the two shoulders, ML separation at crossing is closer to 20 – 50 cm in lab-based studies. In our study, average ML separation at crossing was 189.3 ± 58.9 cm (or 144.3 ± 58.9 cm, accounting for the bounding box). It is possible that pedestrians maintained a greater ML separation in our study because of the COVID-19 pandemic (which occurred during the time when we collected the video footage). Indeed, the pandemic notably increased personal space for people having a conversation (Holt et al., 2022). Alternatively, pedestrians may naturally adopt a strategy of maintaining a larger ML separation in open, outdoor spaces. We believe this latter explanation is more likely given the summer months and outdoor atmosphere when and where data collection occurred. Indeed, the path where pedestrians walked was approximately 3.5 m wide, excluding the dirt/grass surrounding it, which is wider than most lab spaces and affords greater ML separation at the time of crossing.

Crowds can interfere with one’s ability to walk toward a goal and challenge collision avoidance behaviour. We found that crowd size between approaching pedestrians at 8 m apart predicted path deviation, such that pedestrians were less likely to deviate with greater crowd size for a given ML separation. Virtual reality and simulation studies also demonstrate that crowd characteristics can impact pedestrian behaviour (Bruneau et al. 2015; Koilias et al. 2020; Moussaïd et al. 2011; Murakami et al. 2019; Nelson et al. 2019). For instance, people are more likely to deviate when encountering a group of pedestrians with small interpersonal distance (i.e., greater crowd density) or when the group’s motion is diagonal or orthogonal to their own (Bruneau et al. 2015; Koilias et al. 2019). What could explain our findings? Each pedestrian has the possibility of changing their path (independent of others) and pose a threat for a potential collision. The probability of colliding with one of them increases with greater crowd size. Thus, if one deviates from their path too early or too much, they risk the chance of colliding with another pedestrian or having to make multiple (costly) trajectory corrections.

Our results also revealed an association between greater ML separation at the time of crossing when one or both pedestrians were distracted. Looking around, having a conversation with someone you are walking with, or using a cell phone, are distractions that divert attention from the immediate surroundings and thus, have the potential to affect collision avoidance behaviour. This may relate to decreased situational awareness (Feld and Plummer 2019; Lim et al. 2015; Lin and Huang 2017). Indeed, text messages can increase obstacle detection time (Souza Silva et al. 2019). Furthermore, cell phone use during walking is associated with increased lateral deviation (Lamberg and Muratori 2012). Similarly, Gerin-Lajoie et al. (2006) found that auditory distraction led to an increase in personal space during collision avoidance. In our study, maintaining a wider margin between an approaching pedestrian may be a result of the distracted pedestrian deviating away (Lamberg and Muratori 2012), or the non-distracted pedestrian deviating due to the uncertainty of how the approaching distracted pedestrian would behave (Murakami et al. 2022).

Aside from distractions, constraints to one’s actions can also affect one’s avoidance behaviours. We observed that the presence of a mobility constraint associated with smaller ML separation at the time of crossing. Why might this occur? It is possible that the mobility constraint makes it difficult for the person to deviate; consider walking a bicycle along a path or pushing a bulky stroller. The other pedestrian may recognize this and consider it less uncertain as to whether the mobility-constrained pedestrian can/will deviate. Thus, the pedestrian may be more comfortable with a smaller ML separation. We are unaware of any lab-based study that has examined the effect of a pedestrian with a mobility constraint (e.g., pushing a stroller) on collision avoidance behaviour. Thus, future studies should explore these situations in greater detail.

Although our study offers valuable insights into collision avoidance behaviour in a natural environment, it has limitations. For instance, our data is based on one-on-one interactions, which may not fully reflect the complexity of pedestrian interactions. In addition, our data is from a single location with a linear walking path. Thus, our results may not generalize to different environments. We also had only one interaction per pair of pedestrians, and we could not control the situation, nor could we replicate any situation multiple times. Furthermore, due to a limited field of view of our video camera, we were unable to capture all pedestrian interactions; some interactions occurred too close to the borders of the scene image to extract enough trajectory information. Occasionally, the pedestrian detection algorithm also failed to detect a pedestrian for the entire duration of them being in view. Another limitation is the lack of personal data on the pedestrians. Thus, we were unable to determine how sex or body size, for example, influence behaviour.

Besides providing important fundamental insight into collision avoidance behaviour, our findings may facilitate urban planning and the advancement of autonomous robots. For instance, one can improve the design and safety of public walkways and sidewalks by considering the factors that affect pedestrian walking behaviour. For robots that must navigate through crowded public spaces, scientists and engineers could incorporate algorithms for maintaining adequate personal space and that account for the impact of distractions and pedestrian mobility constraints. Embedding these human behavioural models could enable more seamless integration into pedestrian flows, ensuring that robots can navigate complex social spaces without disrupting the natural movement patterns of humans.

In conclusion, our findings demonstrate that collision avoidance behaviours observed in a lab setting are similar to those observed in a real-world setting (even if the specific values of the measures differ). This is particularly important given that we assessed multiple factors simultaneously as opposed to in isolation like most lab-based research. Crucially, observing people interact in a natural environment can help inform future research questions and lab-based study designs to better understand collision avoidance behaviours.

## Acknowledgements

The authors wish to thank all students involved in helping to characterize interaction details.

## Notes

### Competing Interest Statement

The authors have declared no competing interest.

